## Supplemental Methods for "Intracellular diversity of WNV within circulating avian peripheral blood mononuclear cells reveals host-dependent patterns of polyinfection"

**Supplementary Methods 1.**

**Establishing an Amplicon Targeted Sequencing Approach**

To overcome sequencing errors and PCR-mediated recombination that might introduce biases (1-4), we adjusted a library preparation strategy established by others (5). This strategy offers the inclusion of a stretch of degenerate nucleotides included into the cDNA synthesis primer that enables “tagging” of the input cDNA template before proceeding on to PCR amplification steps. This stretch of nucleotides served as an identifier sequence and was previously referred to as “Primer ID”(5).

Since the focus of this study was to identify intra-host viral genetic diversity and we specifically aimed at sequencing single cells carrying a plethora of viral genomes, it was important to be able to identify low frequency genomes and to have as low sequencing bias as possible The inclusion of the Primer ID sequence allowed each original template copy to be tagged with an identifier sequence. Finding multiple identical Primer ID sequences during the analysis process of the sequencing products indicated that genomes tagged with the same Primer ID originated from a single original input template. These were subsequently collapsed to create a consensus sequence for each original template.

We adapted the Primer ID approach to the Illumina MiSeq platform and amplified a 335 bp long fragment within the WNV genome that included the barcoded region within the NS4b segment (Supplementary figure 1, table of Illumina barcodes used for this study, excel sheet table-Illumina barcodes used for sample multiplexing (6). Library preparation is further detailed in the Experimental procedures section. Using the Primer ID method we were able to generate libraries from an input of as little as 50 viral genomes (Supplementary Methods Figure 1).

**Supplementary Methods Figure 1. (A)** Table of illumina barcodes. (**B**) Table of primers for illumina sample multiplexing and primer ID. (**C**) Determination of limit of Primer ID detection.
