## Supplementary figures and images for "Intracellular diversity of WNV within circulating avian peripheral blood mononuclear cells reveals host-dependent patterns of polyinfection"

### Supplemental Methods Figure

#### Slide 1
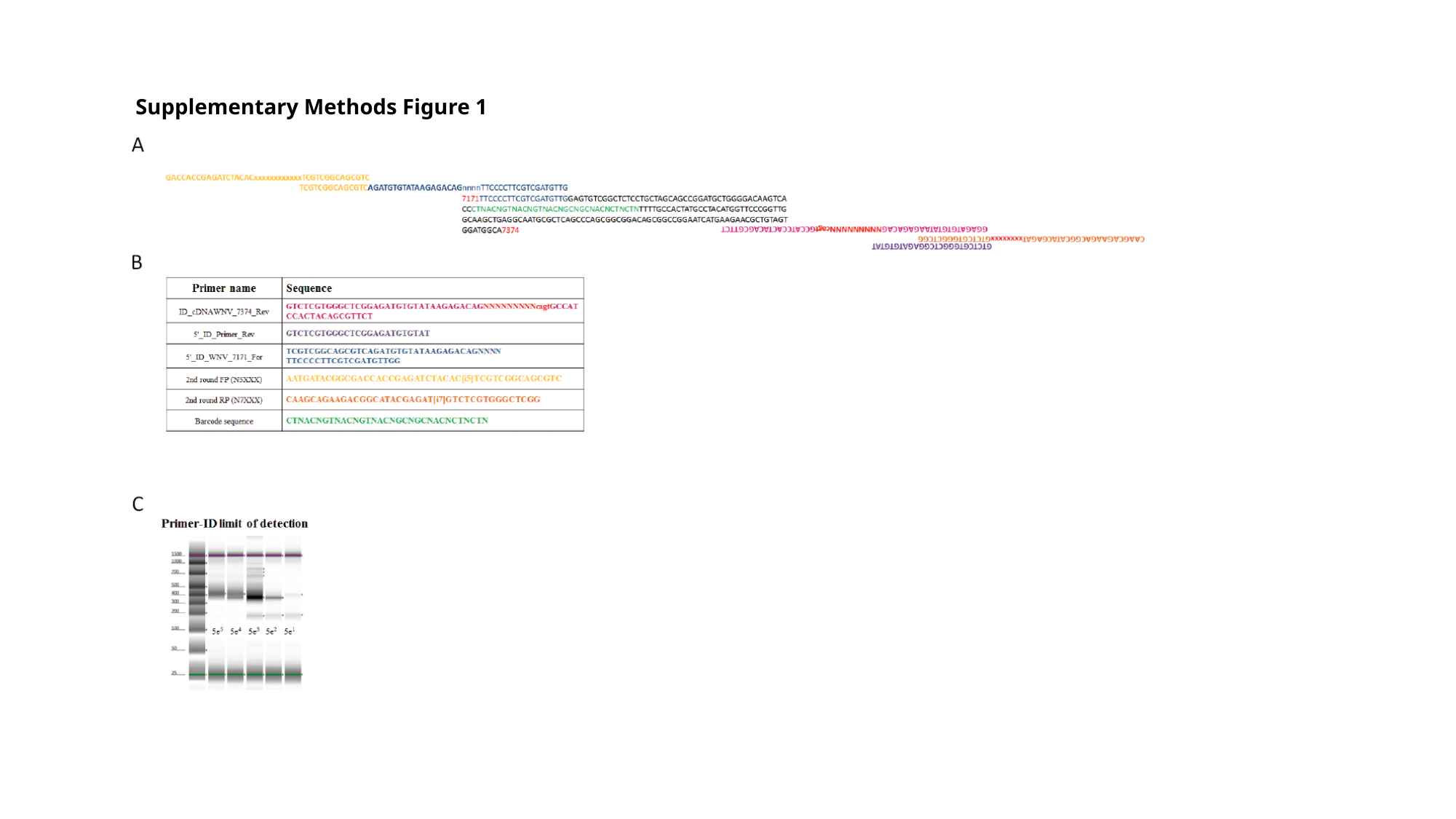

Supplementary Methods Figure 1
